# Evaluation of the virucidal efficacy of Klaran UVC LEDs against surface dried norovirus

**DOI:** 10.1101/2021.04.28.441855

**Authors:** Richard M. Mariita, Amy C. Wilson Miller, Rajul V. Randive

## Abstract

Human norovirus (HuNoV) is a highly contagious pathogenic virus that is transmitted through contaminated food, water, high-touch surfaces and aerosols. Globally, there are an estimated 685 million infections annually due to norovirus, among them 200 million children under the age of 5, causing approximately 50,000 child deaths per year and costing an estimated $60 billion annually in healthcare. In the USA, HuNoV is responsible for 19-21 million illnesses, with an average of 570-800 deaths per year. HuNoV is especially pernicious because it requires less than 100 viral particles to cause an infection, and there are few effective disinfectants. It is believed that Ultraviolet Subtype C (UVC) irradiation might prove to be an effective disinfectant. This study seeks to determine the inactivation profile of UVC against norovirus using a Klaran UVC Light-emitting diode (LED) array product number KL265-50V-SM-WD, emitting radiation at 269 nm peak wavelength and a measured fluence of 1.25 mW/cm^2^ at a 7 cm source-surface distance. Since the HuNoV cannot currently be propagated in cell cultures, the study utilized feline calicivirus (FCV), a recommended surrogate as challenge organism. The test followed Modified ASTM E2197. Assessment of virus inactivation was performed using plaque assay method, with Crystal Violet as a staining agent to enhance plaque visualization. Within 18 seconds of exposure time at a UVC irradiance of 1.25mW/cm^2^ and a dose of 22.5 mJ/cm^2^, the study obtained 99.9 % virus reduction (3 log reduction value). These results demonstrate that Klaran UVC-LED array (KL265-50V-SM-WD) can provide effective inactivation of HuNoV.

## Introduction

Human norovirus (HuNoV), formerly known as Norwalk virus is an RNA virus belonging to *Caliciviridae* family (1). This is a highly contagious, small, non-enveloped enteric pathogen that is transmitted through person-to-person contact and unsanitary food handling (2), contaminated water and high-touch surfaces (1) and can also be spread via aerosols (3). Because it only requires small inoculum to produce an infection (<100 viral particles), its pathogenicity and the ability to survive in different environments, HuNoV is responsible for substantial comorbidity, especially in health care and community settings (1), such as daycare centers, nursing homes, hospital wards, schools, restaurants, catered events and cruise ships (4).

Globally, there are an estimated 685 million cases of norovirus infections annually, with about 200 million of them being among children under 5 years, leading to an estimated 50,000 child deaths and healthcare cost estimation of $60 billion per year (5). In the USA, HuNoV is associated with 80-90% of the reported outbreaks and is the leading cause of nonbacterial gastroenteritis (4). On average, in the USA, HuNoV causes an average of 570-800 deaths, 56,000-71,000 hospitalizations, 400,000 ER visits, 1.7-1.9 million outpatient visits, and 19-21 million total illnesses per year (6). Outbreaks involve people in high-risk groups, particularly young children under 5 years of age, the elderly above 65 (6), travelers, soldiers and the immunocompromised (4). To compound the problem, presently, HuNoV has a limited number of disinfectants that are effective against them (7).

Study of HuNoV has been hindered by the inability to propagate it in cell cultures (8). Because of that, another virus from the same *Caliciviridae* family, Feline calicivirus (FCV) is frequently used as a surrogate (9), especially in determining virucidal efficacy of disinfectants (7). For RNA viruses, Ultraviolet Subtype C (UVC) induces inactivation that leads to RNA damage (10), principal to loss of viral infectivity (11), and FCV is thought to be a reasonable surrogate for HuNoV’ UVC inactivation profiles. Light-emitting diodes (LEDs) made from semiconductor materials can be used to produce UVC in the range of 200-280 nm (12) that is considered to be germicidal (13). LEDs emitting UVC have been used in agriculture, water and the food industry for microbial inactivation because they have several advantages over conventional sources (12). Such advantages include compact size that makes incorporation easier, nonhazardous (no health hazards due to possibility of mercury contamination), high performance and lifetime (12).

Because of the existing economic and public burden associated with HuNoV, there is need for additional appropriate interventions, including effective inactivation strategies. Here, the study investigated the virucidal activity of UVC LED array (product number KL265-50V-SM-WD) against FCV on magnetic stainless-steel discs. Steel coupons were used because human noroviruses transmission can be via contaminated surfaces (11).

## Methods

A USB4000 photospectrometer (Ocean Optics) was used to confirm the emitted radiation peak wavelength of the UVC LED array. For UVC dose, confirmation was achieved using X1 handheld optometer (Gigahertz-Optik). The UVC LED array tested in this study was product number KL265-50V-SM-WD, which is rated between 70-80 mW at 500 mA and was driven at 350 mA, yielding an expected 56-64 mW at beginning of life. Eagle's modified medium with 2% Fetal Bovine Serum was used as test solution. The virus was applied on 20 mm diameter magnetic stainless-steel discs, spread and dried at room temperature prior to exposure to UVC. The sample was put in the center of the box, directly opposite and aligned to the light source, where maximum intensity is found. Distance between the LED and microbe was fixed at 7 cm.

The inactivation experiments were performed in duplicate per irradiation period using Feline calicivirus (FCV) spread on stainless steel magnetic discs following a modified ASTM E2197: Standard Quantitative Disk Carrier Test Method (14). Specifically, this was done by applying 32 µL of standardized FCV on each disc, spreading to within 1 mm of the edges, and then air drying at room temperature. The stainless steel magnetic discs were then irradiated for 12, 18 and 22 seconds prior to suspension in 10 mL modified SCDLP (Soybean, Casein Digest Agar with Lecithin and Polysorbate) medium for virus recovery.

To quanitify viable viral particles, dilution plate method was followed, with Crandall-Rees Feline

Kidney (CRFK) cell lines being used for growth (15). Viral particles were determined using Crystal Violet for staining to enable visualization of plaques for counting, giving Plaque Forming Units (PFU) (16). Mean PFUs between controls and irradiated samples were then used to calculate reduction following the formula:

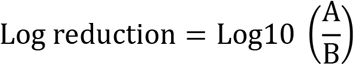

Where:

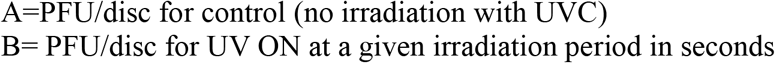

## Results and discussion

The confirmed peak wavelength of the UVC array was 269 nm, and at 7 cm height (Figure 1), it obtained an intensity of 1.25 mW/cm^2^. At 12 seconds and a dose of 15 mJ/cm^2^, the LED array obtained 2.70 log reduction of viable FCV virus (Table 1). At 18 seconds or more (22.5 mJ/cm^2^ or higher dose), >3 log reduction of viable FCV virus was obtained. No inactivation differences at 18 and 22 seconds (22.5 mJ/cm2 and 27.5 mJ/cm2) of irradiation were revealed (Table 1).

**Figure 1:**
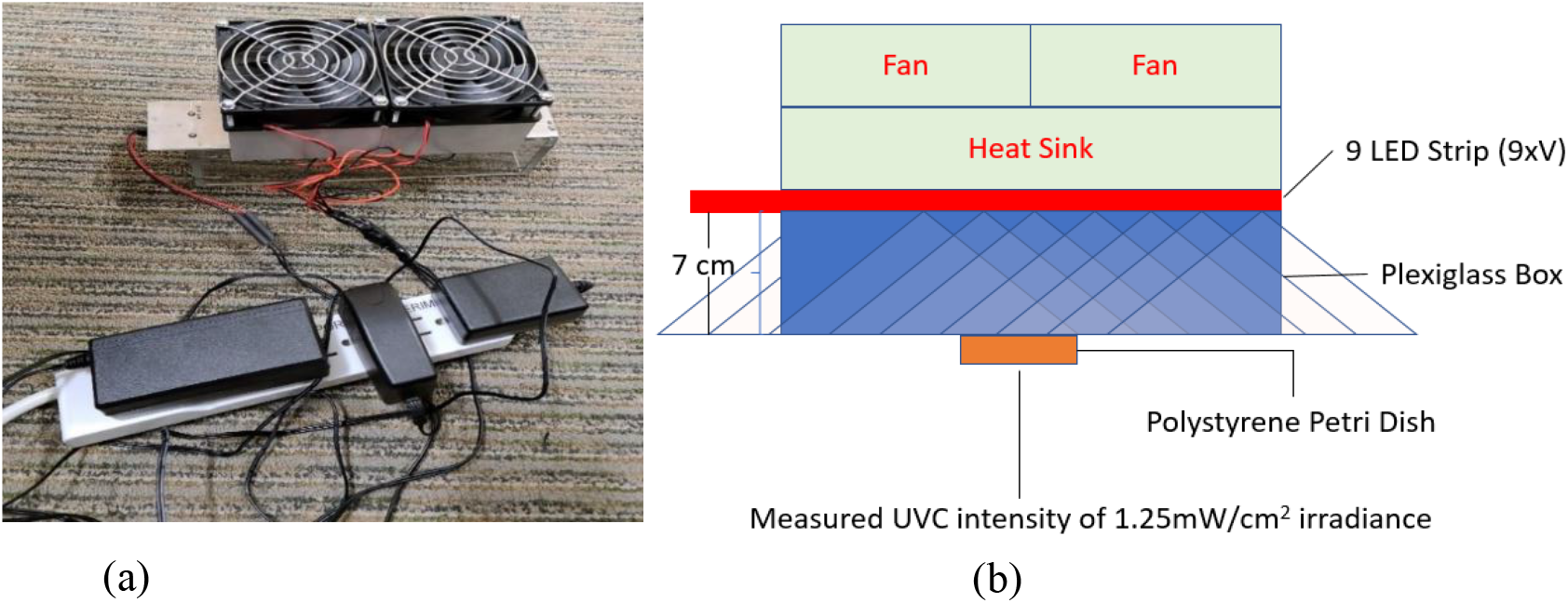
(a) The UVC array was driven at 350 mA during the norovirus inactivation study. (b) The array had two fans and a heat sink for thermal management and was tested at 7 cm height.

**Table 1:**
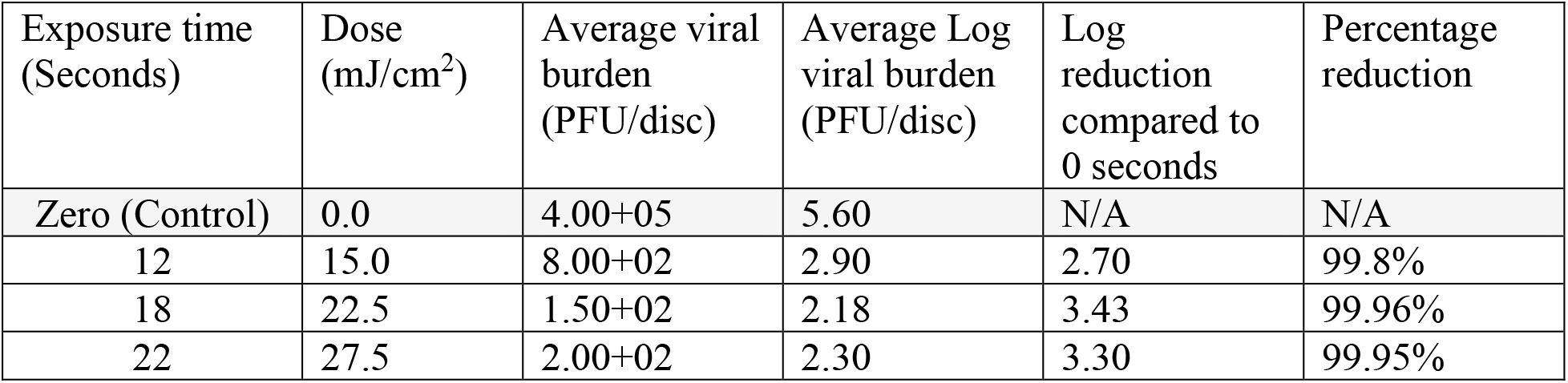
Inactivation performance of an array (product number KL265-50V-SM-WD) emitting UVC radiation at 269 nm peak wavelength with intensity of 1.25 mW/cm^2^ revealed no inactivation differences at 18 and 22 seconds of irradiation. A dose of 22.5 mJ/cm2 was enough to obtain >3log reduction.

| Exposure time (Seconds) | Dose (mJ/cm<sup>2</sup>) | Average viral burden (PFU/disc) | Average Log viral burden (PFU/disc) | Log reduction compared to 0 seconds | Percentage reduction |
| --- | --- | --- | --- | --- | --- |
| Zero (Control) | 0.0 | 4.00+05 | 5.60 | N/A | N/A |
| 12 | 15.0 | 8.00+02 | 2.90 | 2.70 | 99.8% |
| 18 | 22.5 | 1.50+02 | 2.18 | 3.43 | 99.96% |
| 22 | 27.5 | 2.00+02 | 2.30 | 3.30 | 99.95% |

These results are specific to 269 nm against FCV, the commonly used HuNoV model organism. Although spectral sensitivity of HuNoV has not been investigated, it should be expected that against viral pathogens, different wavelengths will perform differently, even with the same dose (17). In Coronaviruses for instance, a study by Gerchman et. al., (17), has demonstrated that UVC LEDs emiting radiation at peak wavelength between 267 nm and 279 nm were more effective in inactivation. Similar approach, however, should be utilized to confirm performance against HuNoV so as to determine sensitivities.

The findings from the current study demonstrate potential application of product KL265-50V-SM-WD UVC arrays in cruise ships and resorts, especially living and dining quarters, where there are high risks in HuNoV acquisition which can lead to disease outbreak (18). With necessary radiation safety considerations, results can further be applied in other areas of close living quarters or with shared dining facilities to help disrupt viral transmission (1). Such areas include those that have reported norovirus outbreaks such as schools (19), military training centers and fields of operation (20), healthcare facilities (21), restaurant and catering industry (22) as well as municipal and industrial water systems (23).

## Conclusion

There are a limited number of virucidal agents known to be presently effective against norovirus (7). There is therefore a need to develop alternative inactivation strategies that are effective against human norovirus, and in particular, strategies that are rapid and chemical-free. In this study, 22.5 mJ/cm^2^ UVC dose was found to be sufficient in order to achieve >3 log reduction against a virus, which is otherwise known to be resistant against most disinfectants (7). The exhibited virucidal activity against FCV suspensions air dried on stainless steel magnetic discs is promising because surfaces are vectors of HuNoV transmission during outbreaks (24). The use of UVC LEDs thus promises a reduction of virus transmission during outbreaks.

These findings demonstrated that UVC LEDs could serve as an effective and rapid tool in the fight against human norovirus by preventing spread via fomites.

## Data availability

Original laboratory report with data is available upon request.

## Acknowledgments

Authors thank Dr. Kevin Kahn, James Davis and Michelle Lottridge for their help with review of manuscript. Authors also thank the ResInnova Labs (Silver Spring, MD, USA), an International Antimicrobial Council (IAC) certified laboratory for utilizing their virology capabilities in the Norovirus experiments for Crystal IS.

## Ethical standards

The manuscript does not contain data obtained through clinical studies or patients.

## References

1. Robilotti E, Deresinski S, Pinsky BA. Norovirus. Clin Microbiol Rev. 2015 Jan 1;28(1):134.

2. Hassard F, Sharp JH, Taft H, LeVay L, Harris JP, McDonald JE, et al. Critical Review on the Public Health Impact of Norovirus Contamination in Shellfish and the Environment: A UK Perspective. Food Environ Virol. 2017/02/07 ed. 2017 Jun;9(2):123–41.

3. Jones RM, Brosseau LM. Aerosol Transmission of Infectious Disease. Journal of Occupational and Environmental Medicine [Internet]. 2015;57(5). Available from: https://journals.lww.com/joem/Fulltext/2015/05000/Aerosol_Transmission_of_Infectious_Disease.4.aspx

4. Glass RI, Parashar UD, Estes MK. Norovirus gastroenteritis. N Engl J Med. 2009 Oct 29;361(18):1776–85.

5. CDC. Norovirus Worldwide [Internet]. 2021. Available from: https://www.cdc.gov/norovirus/trends-outbreaks/worldwide.html

6. Hall AJ, Lopman BA, Payne DC, Patel MM, Gastañaduy PA, Vinjé J, et al. Norovirus disease in the United States. Emerg Infect Dis. 2013 Aug;19(8):1198–205.

7. Whitehead K, McCue KA. Virucidal efficacy of disinfectant actives against feline calicivirus, a surrogate for norovirus, in a short contact time. Am J Infect Control. 2010 Feb;38(1):26–30.

8. Kniel KE. The makings of a good human norovirus surrogate. Current Opinion in Virology. 2014 Feb 1;4:85–90.

9. Kampf G, Grotheer D, Steinmann J. Efficacy of three ethanol-based hand rubs against feline calicivirus, a surrogate virus for norovirus. Journal of Hospital Infection. 2005;60(2):144–9.

10. Beck SE, Rodriguez RA, Hawkins MA, Hargy TM, Larason TC, Linden KG. Comparison of UV-Induced Inactivation and RNA Damage in MS2 Phage across the Germicidal UV Spectrum. Dozois CM, editor. Appl Environ Microbiol. 2016 Mar 1;82(5):1468.

11. Rönnqvist M, Mikkelä A, Tuominen P, Salo S, Maunula L. Ultraviolet Light Inactivation of Murine Norovirus and Human Norovirus GII: PCR May Overestimate the Persistence of Noroviruses Even When Combined with Pre-PCR Treatments. Food and Environmental Virology. 2014 Mar 1;6(1):48–57.

12. Prasad A, Du L, Zubair M, Subedi S, Ullah A, Roopesh MS. Applications of Light-Emitting Diodes (LEDs) in Food Processing and Water Treatment. Food Engineering Reviews. 2020 Sep 1;12(3):268–89.

13. Kebbi Y, Muhammad AI, Sant’Ana AS, do Prado-Silva L, Liu D, Ding T. Recent advances on the application of UV-LED technology for microbial inactivation: Progress and mechanism. Comprehensive Reviews in Food Science and Food Safety. 2020 Nov 1;19(6):3501–27.

14. Antimicrobial Testing Methods & Procedures: MB-31-03. https://www.epa.gov/pesticide-analytical-methods/antimicrobial-testing-methods-procedures-mb-31-03. 2014;

15. Whittemore JC, Hawley JR, Jensen WA, Lappin MR. Antibodies against Crandell Rees feline kidney (CRFK) cell line antigens, alpha-enolase, and annexin A2 in vaccinated and CRFK hyperinoculated cats. J Vet Intern Med. 2010 Apr;24(2):306–13.

16. Mendoza EJ, Manguiat K, Wood H, Drebot M. Two Detailed Plaque Assay Protocols for the Quantification of Infectious SARS-CoV-2. Current Protocols in Microbiology. 2020 Jun 1;57(1):cpmc105.

17. Gerchman Y, Mamane H, Friedman N, Mandelboim M. UV-LED disinfection of Coronavirus: Wavelength effect. Journal of Photochemistry and Photobiology B: Biology. 2020 Nov 1;212:112044.

18. Wikswo ME, Cortes J, Hall AJ, Vaughan G, Howard C, Gregoricus N, et al. Disease transmission and passenger behaviors during a high morbidity Norovirus outbreak on a cruise ship, January 2009. Clinical infectious diseases. 2011;52(9):1116–22.

19. Prevention (CDC C for DC and. Norovirus outbreak in an elementary school--District of Columbia, February 2007. MMWR Morbidity and mortality weekly report. 2008;56(51– 52):1340–3.

20. McCarthy M, Estes MK, Hyams KC. Norwalk-like virus infection in military forces: epidemic potential, sporadic disease, and the future direction of prevention and control efforts. The Journal of infectious diseases. 2000;181(Supplement_2):S387–91.

21. Zingg W, Colombo C, Jucker T, Bossart W, Ruef C. Impact of an outbreak of norovirus infection on hospital resources. Infection Control & Hospital Epidemiology. 2005;26(3):263–7.

22. Kuo H-W, Schmid D, Jelovcan S, Pichler A-M, Magnet E, Reichart S, et al. A foodborne outbreak due to norovirus in Austria, 2007. Journal of food protection. 2009;72(1):193–6.

23. Kaplan JE, Goodman RA, Schonberger LB, Lippy EC, Gary GW. Gastroenteritis due to Norwalk virus: an outbreak associated with a municipal water system. The Journal of infectious diseases. 1982;146(2):190–7.

24. Jimenez L, Chiang M. Virucidal activity of a quaternary ammonium compound disinfectant against feline calicivirus: a surrogate for norovirus. Am J Infect Control. 2006 Jun;34(5):269–73.

